# Mechanisms of transmission ratio distortion at hybrid sterility loci within and between *Mimulus* species

**DOI:** 10.1101/150854

**Authors:** Rachel E. Kerwin, Andrea L. Sweigart

## Abstract

Hybrid incompatibilities are a common correlate of genomic divergence and a potentially important contributor to reproductive isolation. However, we do not yet have a detailed understanding of how hybrid incompatibility loci function and evolve within their native species, or why they are dysfunctional in hybrids. Here, we explore these issues for a well-studied, two-locus hybrid incompatibility between *hybrid male sterility 1* (*hms1*) and *hybrid male sterility 2* (*hms2*) in the closely related yellow monkeyflower species *Mimulus guttatus* and *M. nasutus*. By performing reciprocal backcrosses with introgression lines, we find evidence for gametic expression of the *hms1-hms2* incompatibility. Surprisingly, however, hybrid transmission ratios at *hms1* do not reflect this incompatibility, suggesting additional mechanisms counteract the effects of gametic sterility. Indeed, our backcross experiment shows hybrid transmission bias toward *M. guttatus* through both pollen and ovules, an effect that is particularly strong when *hms2* is homozygous for *M. nasutus* alleles. In contrast, we find little evidence for *hms1* transmission bias in crosses within *M. guttatus*, providing no indication of selfish evolution at this locus. Although we do not yet have sufficient genetic resolution to determine if hybrid sterility and transmission ratio distortion map to the same loci, our preliminary fine-mapping uncovers a genetically independent hybrid lethality system involving at least two loci linked to *hms1*. This fine-scale dissection of transmission ratio distortion at *hms1* and *hms2* provides insight into genomic differentiation between closely related *Mimulus* species and reveals multiple mechanisms of hybrid dysfunction.

## INTRODUCTION

Hybrid incompatibilities are a common outcome of genomic divergence among closely related species. Across diverse taxa, a number of genes for hybrid inviability and sterility have been identified (see Presgraves 2010; Maheshwari and Barbash 2011; Sweigart and Willis 2012; Ouyang and Zhang 2013), but we still know very little about how such genes function and initially evolve within their native species. One possibility is that the initial mutations are selectively neutral and become fixed by random genetic drift. Alternatively, the mutations might increase in frequency because they benefit the native species for reasons that are incidental to their role in reproductive isolation – by promoting ecological adaptation, for example (Schluter and Conte 2009). Yet another possibility is that hybrid incompatibilities arise through recurrent bouts of intragenomic conflict within species (Frank 1991; Hurst and Pomiankowski 1991). In this last scenario, selfish genetic elements (*e.g*., transposons, meiotic drivers, gamete killers) manipulate host reproduction to bias their own transmission. Because these actions are often detrimental to host fitness, there is then selective pressure for compensatory mutations or suppressors to neutralize the effects of selfish evolution (Burt and Trivers 2006).

The idea that intragenomic conflict involving segregation distorters might be a major source of hybrid incompatibilities has resurged in recent years (Johnson 2010; McDermott and Noor 2010; Presgraves 2010; Crespi and Nosil 2013), largely due to influential studies in *Drosophila* that have mapped hybrid segregation distortion and hybrid sterility to the same genomic locations (Tao et al. 2001; Phadnis and Orr 2009b; Zhang et al. 2015). In plants, too, classic and recent crossing studies have revealed “gamete killers” that affect both transmission ratios and fertility; at these loci, one parental allele causes the abortion of gametes carrying the other allele (*e.g*., tobacco: (Cameron and Moav 1957), wheat: (Loegering and Sears 1963), tomato: (Rick 1966), rice: (Sano 1990; Long et al. 2008; Yang et al. 2012), Arabidopsis: (Simon et al. 2016)). Although suggestive of a causal link between selfish genetic elements and hybrid incompatibilities, few studies have proven a history of segregation distortion *within* species. Thus, in most cases, an alternative possibility is that segregation distortion acts exclusively in hybrid genetic backgrounds, and is a consequence rather than a cause of the incompatibility.

In seed plants, hybrid incompatibilities can act in either the diploid sporophyte or the haploid gametophyte, two stages of the life cycle that are controlled by different sets of genes and subject to distinct evolutionary forces (Walbot and Evans 2003; Gossmann et al. 2014; Gossmann et al. 2016). Unlike in animal systems, which have very little haploid gene expression in sperm or egg cells (Braun et al. 1989; Barreau et al. 2008), thousands of genes are expressed in plant gametophytes (*i.e*., pollen and embryo sacs in angiosperms) (Wuest et al. 2010; Rutley and Twell 2015). As a result, hybrid sterility in plants can be caused by genetic incompatibilities that affect the haploid gametophytes or the diploid sporophytic tissues surrounding the gametes (*e.g*., tapetum for pollen, ovule cells for the embryo sac). Of these two possibilities, the former appears to be much more common among the ~50 hybrid sterility loci that have been identified between subspecies of Asian cultivated rice, *Oryza sativa ssp. japonica* and *O. sativa ssp. indica* (Morishima et al. 1991; Ouyang and Zhang 2013). A large number of gametic incompatibilities have also been shown to contribute to transmission ratio distortion in crosses between populations of *Arabidopsis lyrata* (Leppala et al. 2013). This bias toward gametic incompatibilities might be due to differences in the number of mutations that affect the two classes of hybrid sterility and/or to the fact that recessive alleles are exposed in the haploid gametophyte (similar to genes on heteromorphic sex chromosomes). Additionally, rates of evolution might be accelerated for gametophytic genes due to sex-specific selection (Gossmann et al. 2014). It is also possible that intragenomic conflict is more common in the gametophyte; any selfish genetic element that can disable gametes carrying the alternative allele will have a direct impact on its own transmission.

Of the handful of plant hybrid sterility genes that have been cloned, all are in rice, most are gametic, and many appear to have evolved via neutral processes. The two most straightforward examples involve pollen defects caused by loss-of-function alleles at duplicate genes (Mizuta et al. 2010; Yamagata et al. 2010), consistent with a model of divergent resolution via degenerative mutations and genetic drift (Werth and Windham 1991; Lynch and Force 2000). The remaining six cases all involve gamete killers (Long et al. 2008; Kubo et al. 2011; Yang et al. 2012; Kubo et al. 2016a; Kubo et al. 2016b; Yu et al. 2016), which might be taken as evidence for pervasive selfish evolution within rice species. However, molecular characterization of these hybrid sterility systems has provided little support for this scenario. For example, the *S5* locus causes female sterility in *japonica*-*indica* hybrids when gametes carry an incompatible combination of “killer” and “protector” alleles at three, tightly linked genes (Yang et al. 2012). The two domesticated subspecies carry null alleles in distinct components of the killer-protector system. Because both derived haplotypes are perfectly compatible with the ancestral haplotype, it seems unlikely that they entailed fitness costs. Although it is conceivable that intragenomic conflict played a role in the initial formation of the *S5* haplotype (*i.e*., the ancestral killer/protector combination might represent a resolved conflict), it does not seem to be the cause of the current reproductive barrier between *japonica* and *indica*. Similarly, at the *Sa* locus, which causes *japonica*-*indica* hybrid male sterility, patterns of molecular variation and the prevalence of “neutral” alleles that are compatible in all crosses suggest that hybrid dysfunction may have evolved unopposed by natural selection (Long et al. 2008; Sweigart and Willis 2012). A key feature of these gamete killers is that they are caused by two or more tightly linked, epistatic genes (Long et al. 2008; Yang et al. 2012; Kubo 2013; Kubo et al. 2016a). Adding to the complexity, some of them require additional, unlinked loci that act sporophytically (Kubo et al. 2011; Kubo et al. 2016a; Kubo et al. 2016b). Taken together, these studies suggest that hybrid sterility in rice is polygenic and might evolve without significant fitness costs within species. However, it is not yet clear if these themes are generalizable to other plant systems.

Here we investigate patterns of transmission ratio distortion associated with a two-locus hybrid sterility system between the closely related monkeyflower species, *Mimulus guttatus* and *M. nasutus*. Previously, we fine-mapped the two incompatibility loci – *hybrid male sterility 1* (*hms1*) and *hybrid male sterility 2* (*hms2*) – to small nuclear genomic regions of ~60 kb each on chromosomes 6 and 13 (Sweigart and Flagel 2015). We also discovered evidence that the *hms1* incompatibility allele is involved in a partial selective sweep within a single population of *M. guttatus*, but the underlying cause of the sweep is unknown (Sweigart and Flagel 2015). Additionally, because the *hms1* sterility allele is embedded in a nearly invariant, 320-kb haplotype, it is not yet clear whether *hms1* or a linked locus is the target of the sweep. This polymorphic hybrid sterility system provides a unique opportunity to test directly whether selfish evolution within species can lead to incompatibilities between species.

Previously, in crosses between *M. guttatus* and *M. nasutus*, we observed transmission ratio distortion (TRD) of genotypes at both *hms1* and *hms2* (Sweigart et al. 2006; Sweigart and Flagel 2015), but the causes have remained unexplored. Additionally, these previous studies did not test directly whether the *hms1-hms2* incompatibility acts in the gametophyte or sporophyte, although patterns of F_2_ hybrid sterility seemed to suggest the latter. Results from these studies suggested that the incompatibility acts in the diploid sporophyte with the *M. guttatus* allele at *hms1* acting dominantly in combination with recessive *M. nasutus* alleles at *hms2* to cause nearly complete male sterility and partial female sterility (Sweigart et al. 2006). Consistent with this genetic model, pollen viability is ~20% in F_2_ hybrids that are heterozygous for *hms1* and homozygous for *M. nasutus* alleles at *hms2* (*hms1*_GN_; *hms2*_NN_), much lower than the 50% expected for a strictly gametic hybrid incompatibility (with *hms1*_G_; *hms2*_N_ causing dysfunction). Moreover, because a gametic hybrid incompatibility should cause transmission bias at *both* interacting loci, we would expect a deficit of *M. guttatus* alleles at *hms1* equal to that of *M. nasutus* alleles at *hms2*. Although F_2_ hybrids do indeed show a deficit of *M. nasutus* alleles at *hms2*, allelic transmission at *hms1* follows the Mendelian expectation (Sweigart et al. 2006).

In the current study, we used introgression lines (ILs) and a reciprocal backcross design to distinguish among at least four possibilities for TRD in genomic regions linked to *hms1* and *hms2:* 1) distortion through male gametes due to pollen competition and/or pollen sterility, 2) distortion through female gametes due to female meiotic drive (*e.g*., (Fishman and Saunders 2008) and/or ovule sterility, 3) TRD through both male and female gametes due to an incompatibility that affects both gametophytes (*e.g*., (Kubo et al. 2016a), and 4) distortion caused by selection against zygotes. In a series of crossing experiments, we investigated the mechanism of TRD at *hms1* and *hms2* and addressed the following specific questions. Is hybrid transmission bias at *hms1* and/or *hms2* a simple byproduct of gametic hybrid sterility? Is there evidence for hybrid transmission bias at these loci independent of gamete sterility? Are hybrid sterility and TRD genetically separable? Does TRD at *hms1* occur within *M. guttatus*? Our results provide insight into the mechanisms of hybrid sterility and transmission distortion, and into the evolutionary dynamics of incompatibility alleles within species.

## METHODS

### Study system and plant lines

The *Mimulus guttatus* species complex is a group of phenotypically diverse wildflowers with abundant natural populations throughout much of western North America. In this study, we focus on *M. guttatus* and *M. nasutus*, two members of the complex that diverged roughly 200,000 years ago (Brandvain et al. 2014). These species occupy a partially overlapping range, and are primarily differentiated by mating system. *M. guttatus* is predominantly outcrossing with showy, insect-pollinated flowers, whereas *M. nasutus* is highly self-fertilizing with reduced flowers. In geographic regions where the two species co-occur, they are partially reproductively isolated by differences in floral morphology, flowering phenology, and pollen-pistil interactions (Diaz and Macnair 1999; Martin and Willis 2007; Fishman et al. 2014). Hybrid incompatibilities are also common, but variable (Vickery 1978; Christie and Macnair 1987; Sweigart et al. 2007; Case and Willis 2008; Martin and Willis 2010). Despite these barriers to interspecific gene flow, sympatric populations display evidence of genome-wide introgression (Sweigart and Willis 2003; Brandvain et al. 2014; Kenney and Sweigart 2016).

Previous work identified two nuclear incompatibility loci – *hybrid male sterility 1* (*hms1*) and *hybrid male sterility 2* (*hms2*) – that cause nearly complete male sterility and partial female sterility in a fraction of F_2_ hybrids between an inbred line of *M. guttatus* from Iron Mountain, Oregon (IM62), and a naturally inbred *M. nasutus* line from Sherar’s Falls, Oregon (SF5) (Sweigart et al. 2006). In 2015, Sweigart and Flagel generated a large SF5-IM62 F_2_ mapping population (*N* = 5487) to fine map *hms1* and *hms2* to regions of ~60 kb on chromosome 6 and chromosome 13, respectively. Hybrids carrying at least one incompatible *M. guttatus* allele at *hms1* in combination with two incompatible *M. nasutus* alleles at *hms2* display extreme male sterility and partial female sterility (Sweigart et al. 2006). Furthermore, the *hms1* locus is polymorphic within the Iron Mountain population (Sweigart et al. 2007) and several inbred lines derived from that site are known to carry “compatible” alleles that do not cause hybrid sterility when crossed to *M. nasutus* (Sweigart and Flagel 2015). In experimental crosses to test for TRD at *hms1* within *M. guttatus*, we used a compatible line called IM767. In total, three inbred lines were used in different crossing schemes to test for TRD within and between species (see below). SF5 is compatible at *hms1* and incompatible at *hms2,* IM62 is incompatible at *hms1* and compatible at *hms2*, and IM767 is compatible at *hms1* and *hms2*.

All plants were grown in the greenhouse at the University of Georgia. For all crosses, seeds were planted into 96-cell flats containing Fafard 3B potting mix (Sun Gro Horticulture, Agawam, MA), stratified for 7 days at 4°C, and then placed in a greenhouse with supplemental lights set to 16-hr days. Plants were bottom-watered daily and temperatures were maintained at 24°C during the day and 16°C at night.

### Introgression line (IL) crossing design to investigate mechanisms of transmission ratio distortion between *M. guttatus* and *M. nasutus*

Previously, two reciprocal nearly isogenic line (NIL) populations carrying *M. nasutus* (SF5) or *M. guttatus* (IM62) introgressions in the opposite genetic background were generated (Fishman and Willis 2005). Briefly, a single SF5 x IM62 F_1_ and IM62 x SF5 F_1_ individual each served as the initial seed parent then underwent four generations of backcrossing to create a BN_4_ NIL population (SF5 x IM62 F_1_, *M. <u>n</u>asutus* recurrent parent) and BG_4_ NIL population (IM62 x SF5 F_1_, *M. guttatus* recurrent parent). Within the BN_4_ and BG_4_ populations, each NIL carries a unique complement of heterozygous introgressions in a genome that is expected to be 96.875% homozygous for the recurrent parent’s alleles. To determine the genomic locations of the heterozygous introgressed regions, the NILs were genotyped at microsatellite and gene-based markers distributed throughout the genome (L. Fishman, unpublished). We selected three NILs with introgressions spanning *hms1* or *hms2* for further genetic analyses. Against a largely *M. guttatus* background, the BG_4_.476 NIL is heterozygous for an introgression that includes *hms1* and ~78% of the physical distance along chromosome 6. The BG_4_.149 line is heterozygous for an introgression that spans ~71% of chromosome 13 and includes *hms2*.

Against a *M. nasutus* background, the BN_4_.62 line is heterozygous for ~75% of chromosome 13, including *hms2*. In addition to these NILs, we used an *hms1* introgression line, RSB_4_, created after four generations of recurrent selection for hybrid sterility with backcrossing to *M. nasutus,* starting from a sterile SF5-IM62 BC_1_ individual (Sweigart et al. 2006); the heterozygous introgression spans ~50% of chromosome 6.

To characterize TRD between *M. guttatus* and *M. nasutus*, we used a multi-step crossing scheme, starting with the NILs and RSB_4_ (described above), to create a set of lines carrying specific two-locus genotypes at *hms1* and *hms2*. First, to generate introgression lines (ILs) that carry heterozygous alleles at both *hms1* and *hms2* in an otherwise *M. guttatus* or *M. nasutus* genetic background, we crossed BG_4_.476 to BG_4_.149, and BN_4_.62 to RSB_4_. From those progeny, we identified *hms1*-*hms2* double heterozygotes by genotyping markers that flank *hms1* (M8 and M24) and *hms2* (M51 and MgSTS193), as described previously (Sweigart and Flagel 2015). Next, to generate individuals that carry various two-locus combinations at *hms1* and *hms2*, we self-fertilized doubly heterozygous ILs from each genetic background (*i.e*., IL-G and IL-N = *M. guttatus* and *M. nasutus* backgrounds, respectively). These crosses are expected to yield nine different two-locus genotypes each (typical of an F_2_), five of which are heterozygous at *hms1* and/or *hms2* (Figure 1). Surprisingly, one of the relevant IL-N *hms1-hms2* genotypes was not recovered (*hms1*_GG_; *hms2*_GN_, see Figure 1); the *hms1*-introgression could not be made homozygous for *M. guttatus* alleles against an *M. nasutus* genetic background (see Results). We assessed male fertility (*i.e*., pollen viability) for the nine experimental IL genotypes (five for IL-G and four for IL-N) as described previously (Sweigart et al. 2006; Sweigart et al. 2007).

**Figure 1.**
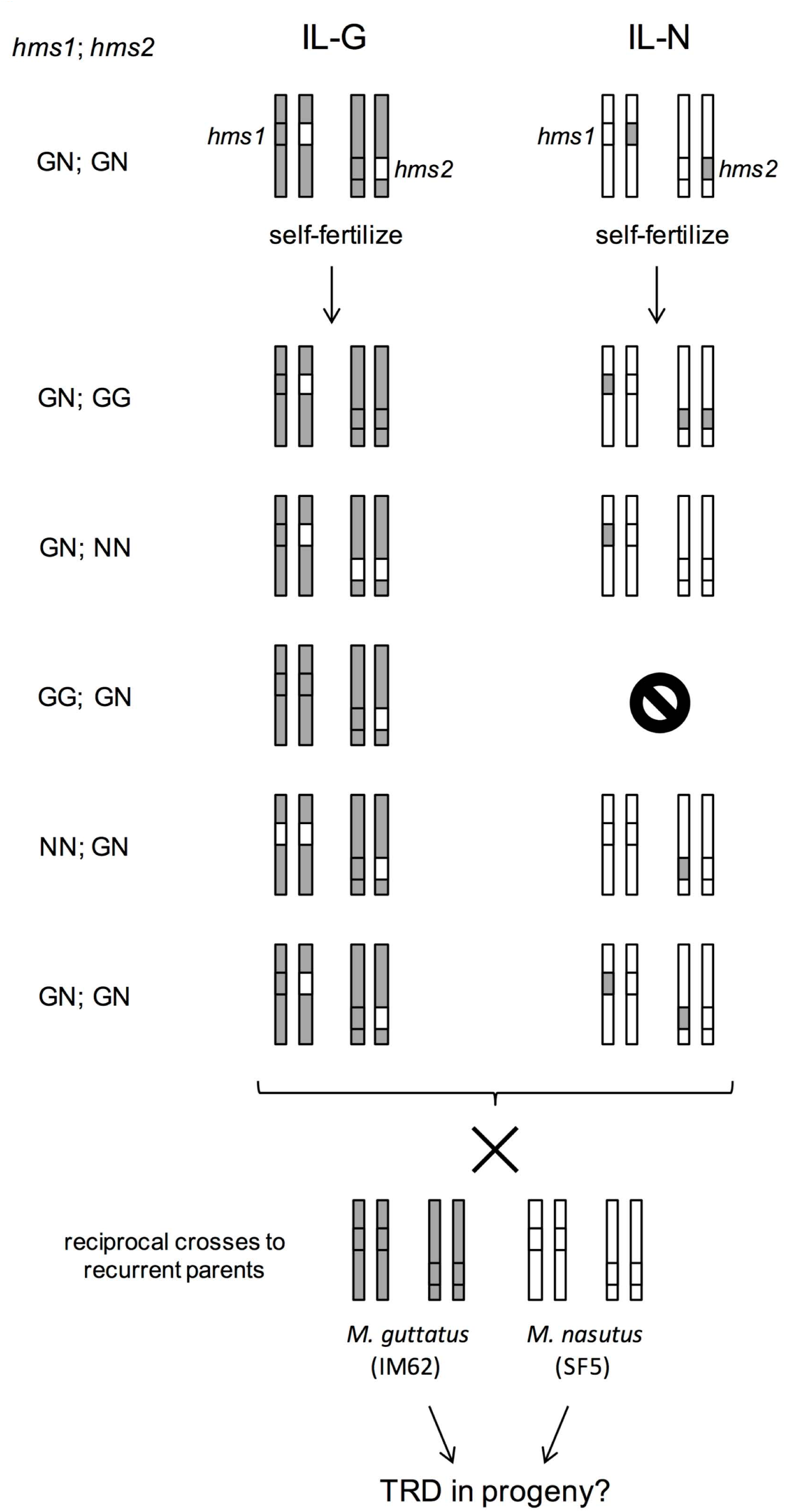
Crossing design for backcross experiment using introgression lines (ILs). For each genotype, two chromosome pairs are shown (one with *hmsl* and one with *hms2*). We constructed two sets of ILs with heterozygous introgressions at both *hmsl* and *hms2;* the IL-G has an *M*. *guttatus* genetic background (grey shading) and the IL-N has an *M. nasutus* genetic background (white). These doubly heterozygous ILs were self-fertilized to generate progeny with two-locus genotypes that are heterozygous at *hmsl* and/or *hms2.* These five progeny types were then reciprocally backcrossed toM *guttatus* and *M. nasutus.* G = M *guttatus* allele (grey); N =*M. nasutus* allele (white).

To test the effect of *hms1* genotype on transmission at *hms2* and vice versa, we reciprocally backcrossed each of the nine ILs to both *M. guttatus* (IM62) and *M. nasutus* (SF5) (Figure 1). Thus, for each IL, we generated four reciprocal backcross populations allowing us to dissect gender-specific transmission ratio distortion. For each IL, two of the backcrosses used the emasculated IL as the seed parent in crosses to IM62 and SF5 lines (i.e., IL-IM62 and IL-SF5) and two used the IL as the pollen parent in crosses to emasculated IM62 and SF5 plants (i.e., IM62-IL, and SF-IL). If *hms* distortion occurs through pollen (due to pollen competition or a gametic incompatibility), we expect TRD in one or both of the backcrosses using the IL as the paternal parent, but not as the maternal parent. If, instead, female meiotic drive and/or a female gametic incompatibility occurs at these *hms* loci, we would expect to see TRD in both backcrosses with the IL as the seed parent, but not with the IL as the pollen parent. Finally, if TRD is caused by the loss of diploid zygotes (or seedlings), it should be apparent in *both* reciprocal crosses to the same recurrent parent (*i.e*., regardless of the gender of the IL). For all crosses, the female parent was emasculated 1-2 days before hand-pollination to prevent self-pollination. Sample sizes for the progeny classes ranged from 33 to 215 individuals (average *N* = 136).

For each *hms* locus, we performed factorial ANOVAs in Jmp Pro 13.0 to examine if genotype ratios were affected by four factors: 1) IL genetic background, 2) IL genotype at the interacting *hms* locus, 3) backcross direction, and 4) identity of the recurrent parent.

### Crossing design to examine transmission ratio distortion within *M. guttatus*

To determine whether transmission ratio distortion at the polymorphic *hms1* incompatibility locus occurs between incompatible and compatible alleles from the Iron Mountain population of *M. guttatus*, we generated reciprocal F_2_ and backcrossed populations with IM62 and IM767. We previously determined that the IM767 inbred line carries a compatible allele at *hms1* (*i.e*., one that does not carry the 320-kb haplotype or cause sterility in combination with SF5 alleles at *hms2*). The IM62 and IM767 inbred lines were intercrossed reciprocally and a single F_1_ hybrid from each was self-fertilized to form reciprocal F_2_ populations (IM62 x IM767: *N* = 267; IM767 x IM62: *N* = 315). To identify putative female- and male-specific sources of TRD, and to distinguish between meiotic/gametic mechanisms versus zygotic selection, we generated reciprocal backcrosses with IM62 and IM767. We used a single F_1_ hybrid (IM62 x 767; maternal parent listed first) to generate four backcross populations to the recurrent parents (F_1_-IM62 BC_1_, IM62-F_1_ BC_1_, F_1_-IM767 BC_1_, IM767-F_1_ BC_1_). Two of these backcrosses used the emasculated F_1_ as the seed parent and two used the F_1_ as the pollen donor in crosses to the emasculated recurrent parents.

We also wanted to examine the effect of *M. nasutus hms2* alleles on patterns of within-*M. guttatus* TRD at *hms1*. We wondered if having *M. nasutus* alleles at *hms2* has the potential to unleash severe distortion at *hms1*, even in an otherwise *M. guttatus* genetic background. To address this question, we intercrossed IM767 with a BG_4_-NIL (BG_4_.275) that is heterozygous for an SF5 introgression spanning ~36% of chromosome 13 including *hms2* (in an IM62 genetic background; Figure S2). We self-fertilized two of the resulting F_1_s to generate F_2_ hybrids segregating for SF5 alleles at *hms2* against an IM62-IM767 F_2_-like genetic background. We then genotyped at *hms*-linked markers (M183 for *hms1* and MgSTS193 for *hms2*) to identify IM62-IM767 *hms1* heterozygotes in combination with three different *hms2* genotypes: 1) IM62 homozygotes, 2) IM767 homozygotes, or 3) SF5 homozygotes. Using each of these three genotypic classes, we performed reciprocal backcrosses to IM767 (Figure S2).

### Assessment of transmission ratio distortion

To examine patterns of TRD at the *hms1* and *hms2* loci, we collected leaf tissue from individual plants and isolated genomic DNA using a rapid extraction protocol (Cheung *et al*. 1993) modified for 96-well format. To infer the *hms1* and *hms2* genotypes of hybrid progeny generated from crosses between IM62 and SF5, we determined genotypes at a multiplexed set of fluorescently labeled markers that flank *hms1* (M8 and M24) and *hms2* (MgSTS193 and M51) following amplification protocols used previously (Sweigart et al. 2006; Sweigart et al. 2007). We excluded individuals with crossovers between either pair of flanking markers; based on expected frequency of double crossovers between flanking markers, genotyping error rates for *hms1* and *hms2* were each < 1%. For experimental crosses involving IM62 and IM767, only one tightly linked marker was used to infer genotype at *hms1* (M183). Based on expected crossovers between *hms1* and M183, the genotyping error rate was < 1%. All fluorescently labeled marker products were run on an ABI 3730 at the University of Georgia Genomics Facility. Genotypes were scored automatically using GeneMarker (SoftGenetics), with additional hand scoring when necessary. We used chi-square tests with two degrees of freedom to determine if *hms*-linked genotypes were significantly distorted.

## RESULTS

### Transmission ratio distortion in *M. nasutus-M. guttatus* F_2_ hybrids

As part of previous efforts to fine-map *Mimulus* hybrid incompatibility loci (Sweigart and Flagel 2015), we generated a large *M. nasutus-M. guttatus* F_2_ hybrid mapping population (*N* = 5487) and genotyped all individuals at gene-based markers flanking *hms1* (M8 and M24) and *hms2* (M51 and MgSTS193). As previously reported (Sweigart et al. 2006; Sweigart and Flagel 2015), we observed significant transmission ratio distortion (TRD) in F_2_ genotypes at both hybrid sterility loci (Table 1). At *hms1*, we observed a significant excess of heterozygotes, but allelic transmission did not differ from the Mendelian expectation. The observed genotype ratios at *hms1* also differed significantly from the expectation given the random union of two gametes with the observed allele frequencies. At *hms2*, we observed an excess of *M. guttatus* homozygotes and a deficit of *M. nasutus* homozygous genotypes, as well as a significant bias toward *M. guttatus* alleles. However, genotype ratios at *hms2* do not differ from what is expected given the observed allele frequencies. Taken together, these patterns suggest TRD at *hms1* might be driven primarily by zygotic selection, whereas *hms2* appears to be influenced primarily by selection among gametes.

**Table 1.**
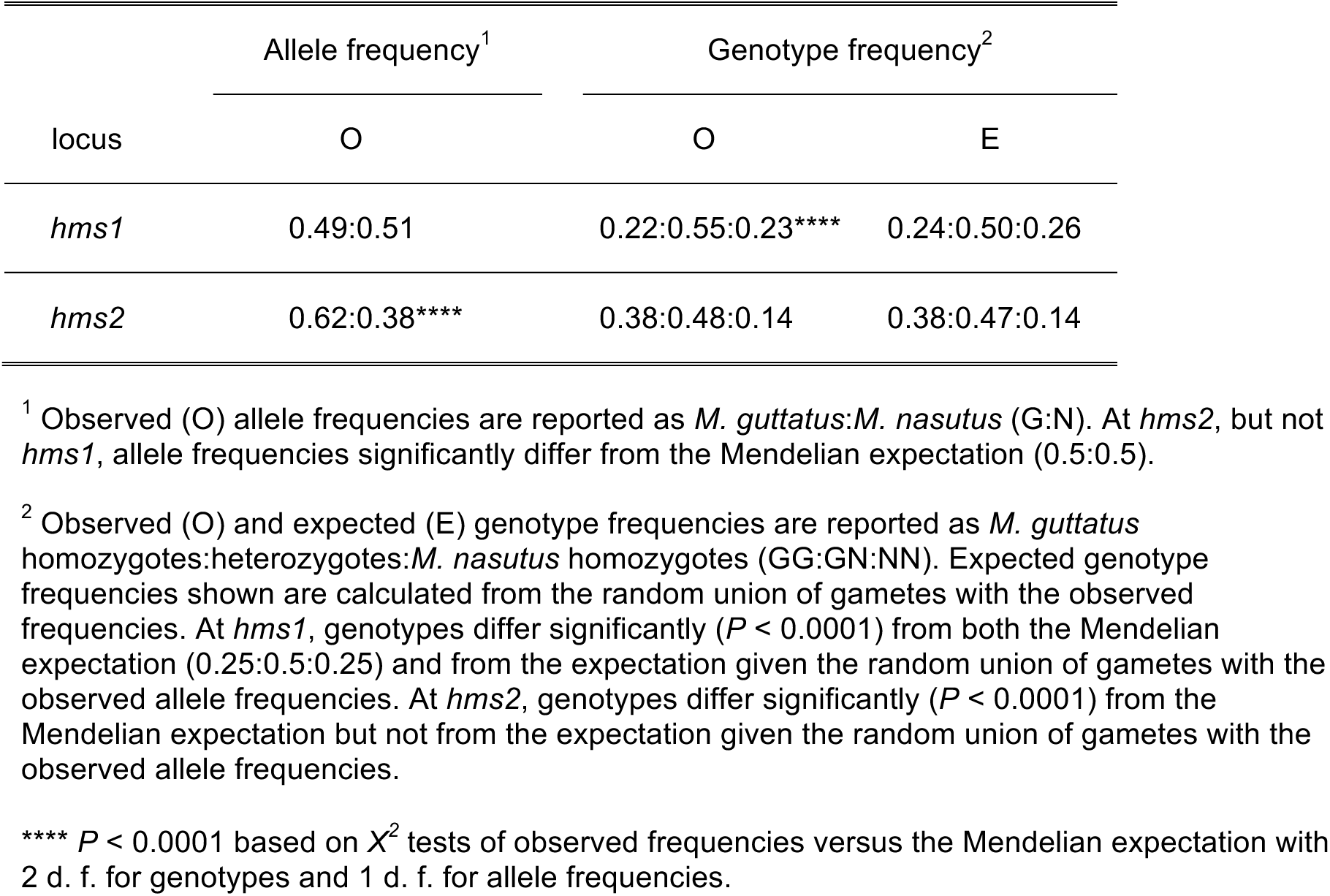
Genotype and allele frequencies at *hmsl* and *hms2* in an *M. nasutus-M. guttatus* F_2_ population (*N* = 5487).

| locus | Allele frequency<sup>1</sup> | Genotype frequency<sup>2</sup> |  |
| --- | --- | --- | --- |
|  | O | O | E |
| *hms1* | 0.49:0.51 | 0.22:0.55:0.23**** | 0.24:0.50:0.26 |
| *hms2* | 0.62:0.38**** | 0.38:0.48:0.14 | 0.38:0.47:0.14 |

<sup>1</sup> Observed (O) allele frequencies are reported as *M. guttatus*:*M. nasutus* (G:N). At *hms2*, but not *hms1*, allele frequencies significantly differ from the Mendelian expectation (0.5:0.5).
<sup>2</sup> Observed (O) and expected (E) genotype frequencies are reported as *M. guttatus* homozygotes:heterozygotes:*M. nasutus* homozygotes (GG:GN:NN). Expected genotype frequencies shown are calculated from the random union of gametes with the observed frequencies. At *hms1*, genotypes differ significantly ( $P < 0.0001$ ) from both the Mendelian expectation (0.25:0.5:0.25) and from the expectation given the random union of gametes with the observed allele frequencies. At *hms2*, genotypes differ significantly ( $P < 0.0001$ ) from the Mendelian expectation but not from the expectation given the random union of gametes with the observed allele frequencies.
\*\*\*\* $P < 0.0001$ based on $\chi^2$ tests of observed frequencies versus the Mendelian expectation with 2 d. f. for genotypes and 1 d. f. for allele frequencies.

When considered together, the two-locus genotypes at *hms1* and *hms2* differ significantly from the Mendelian expectation (*X^2^* = 389.372, d.f. = 8, *P* < 0.0001, *N* = 5487). Although the two-locus genotypes are also significantly different from the expectation given the observed allele frequencies at *hms1* and *hms2* shown in Table 1 (*X^2^*= 71.626, d.f. = 8, *P* <0.0001), the values are much more closely aligned (Table 2). Particularly notable is the deficit of two genotypic classes (*hms1*_GG_; *hms2*_NN_ and *hms1*_NN_; *hms2*_GG_) and the excess of two others (*hms1*_GG_; *hms2*_GG_ and *hms1*_NN_; *hms2*_NN_; Table 2). This pattern of two-locus disequilibrium follows the expectation for gametic action of *hms1-2* sterility (*i.e*., with *hms1*_G_; *hms2*_N_ gametes tending to be sterile). However, the observed F_2_ transmission ratios at *hms1* and *hms2* cannot be entirely explained by *hms1*_G_; *hms2*_N_ gametic sterility (Table S1). This phenomenon, whether acting through one or both parents, would be expected to reduce the transmission of *M. guttatus* alleles at *hms1*, in the same way that it reduces *M. nasutus* alleles at *hms2*. However, there is no indication of allelic transmission bias at *hms1* in the F_2_ hybrids. Taken together, these results suggest that gametic expression of the *hms1-hms2* incompatibility is important, but not the sole contributor, to patterns of transmission ratio distortion in F_2_ hybrids.

**Table 2.** Observed and expected genotype frequencies at *hmsl* and *hms2* in F_2_ hybrids and IL F_2_ hybrids.

| genotype<br>*hms1*; *hms2* | F<sub>2</sub> (5487)<sup>1</sup> |  |  | IL-G F<sub>2</sub> (167)<sup>2</sup> |  | IL-N F<sub>2</sub> (200)<sup>3</sup> |  |  |
| --- | --- | --- | --- | --- | --- | --- | --- | --- |
|  | E: Mendelian | E: obs allele<br>freqs | O | O | E:<br>backcross | O | E:<br>backcross | E: *hms1*<sub>GG</sub> =<br>lethal |
| GG; GG | 0.0625 | 0.093 | 0.099 | 0.066 | 0.107 | 0 | 0.119 | 0 |
| GG; GN | 0.1250 | 0.115 | 0.100 | 0.114 | 0.106 | 0 | 0.069 | 0 |
| GG; NN | 0.0625 | 0.035 | 0.022 | 0.006 | 0.200 | 0 | 0.090 | 0 |
| GN; GG | 0.1250 | 0.191 | 0.208 | 0.174 | 0.176 | 0.185 | 0.193 | 0.241 |
| GN; GN | 0.2500 | 0.236 | 0.268 | 0.234 | 0.249 | 0.300 | 0.249 | 0.310 |
| GN; NN | 0.1250 | 0.073 | 0.071 | 0.054 | 0.078 | 0.075 | 0.058 | 0.073 |
| NN; GG | 0.0625 | 0.098 | 0.070 | 0.102 | 0.072 | 0.085 | 0.077 | 0.096 |
| NN; GN | 0.1250 | 0.121 | 0.117 | 0.180 | 0.133 | 0.225 | 0.151 | 0.188 |
| NN; NN | 0.0625 | 0.037 | 0.047 | 0.072 | 0.061 | 0.130 | 0.0740 | 0.092 |

<sup>1</sup>F<sub>2</sub> genotype counts significantly differ from the Mendelian expectation ( $X^2 = 389.372$ , d.f. = 8, $P < 0.0001$ ) and from what is expected for the random union of gametes given the observed allele frequencies (see Table 1) and independent assortment at *hms1* and *hms2* ( $X^2 = 71.626$ , d.f. = 8, $P < 0.0001$ ).
<sup>2</sup>IL-G F<sub>2</sub> genotype counts significantly differ from the Mendelian expectation ( $X^2 = 18.7910$ , d.f. = 8, $P = 0.0160$ ), but not from what is expected based on allelic transmission in the IL backcrosses (see Table 4, $X^2 = 5.9730$ , d.f. = 8, $P = 0.6502$ ).
<sup>3</sup>IL-N F<sub>2</sub> genotypes significantly differ from the Mendelian expectation ( $X^2 = 86.4090$ , d.f. = 8, $P < 0.0001$ ) and from what is expected based on allelic transmission in the IL backcrosses (see Table 4, $X^2 = 62.0370$ , d.f. = 8, $P < 0.0001$ ), but not from what is expected from the IL backcrosses + *hms1*<sub>GG</sub> homozygote death ( $X^2 = 3.5950$ , d.f. = 5, $P = 0.6090$ ).

### *M. nasutus-M. guttatus* IL crosses reveal multiple causes of F_2_ distortion

To investigate several possible causes of F_2_ transmission ratio distortion at *hms1* and *hms2*, we performed a crossing experiment using the IL-G and IL-Ns. In this crossing design (Figure 1), individuals with one of several possible two-locus *hms1-hms2* genotypes – in each of the IL genetic backgrounds – were crossed reciprocally to *M. guttatus* (IM62) and *M. nasutus* (SF5). By scoring *hms1* and *hms2* genotypes in the progeny of these crosses, we were able to examine the effects of several factors, including parental genotype, genetic background, and cross direction, on transmission ratios at the two hybrid sterility loci. Of the 36 crosses performed, 12 showed significant transmission ratio distortion at *hms1* and/or *hms2* (Table 3; note that two crosses were unsuccessful due to hybrid male sterility). For both *hms1* and *hms2*, parental genotype at one locus has a strong effect on allelic transmission at the other (*hms1* affects *hms2*: *F* = 37.69, *P* < 0.0001; *hms2* affects *hms1*: *F* = 7.80, *P* = 0.004; Figure S1). For *hms2*, cross direction is also important, with stronger TRD occurring through pollen (*F* = 72.33, *P* < 0.0001). Neither the genetic background nor the identity of the recurrent parent significantly affected transmission ratios at *hms1* or *hms2* (results not shown).

**Table 3.** Allelic transmission ratios at *hmsl* and *hms2* in IL-backcross progeny.

| ♀<sup>1</sup> | ♂ | *hms1*; *hms2*<sup>2</sup> | N<sup>3</sup> | *hms1* %G<sup>4</sup> | *hms2* %G<sup>5</sup> |
| --- | --- | --- | --- | --- | --- |
| IL-G | G | GN; GG | 101 | 0.56 |  |
|  |  | GN; NN | 171 | 0.60 |  |
|  |  | GG; GN | 163 |  | 0.53 |
|  |  | NN; GN | 158 |  | 0.47 |
|  |  | GN; GN | 293 | 0.46 | 0.54 |
| IL-G | N | GN; GG | 189 | 0.55 |  |
|  |  | GN; NN | 119 | 0.64* |  |
|  |  | GG; GN | 49 |  | 0.53 |
|  |  | NN; GN | 132 |  | 0.50 |
|  |  | GN; GN | 232 | 0.52 | 0.54 |
| G | IL-G | GN; GG | 382 | 0.55 |  |
|  |  | GN; NN | no seeds | -- |  |
|  |  | GG; GN | 120 |  | 0.86**** |
|  |  | NN; GN | 187 |  | 0.50 |
|  |  | GN; GN | 298 | 0.37*** | 0.67**** |
| N | IL-G | GN; GG | 636 | 0.62**** |  |
|  |  | GN; NN | no seeds | -- |  |
|  |  | GG; GN | 158 |  | 0.90**** |
|  |  | NN; GN | 187 |  | 0.52 |
|  |  | GN; GN | 450 | 0.53 | 0.64**** |
| IL-N | G | GN; GG | 266 | 0.44 |  |
|  |  | GN; NN | 593 | 0.48 |  |
|  |  | GG; GN | n/a |  | -- |
|  |  | NN; GN | 325 |  | 0.55 |
|  |  | GN; GN | 354 | 0.42* | 0.59* |
| IL-N | N | GN; GG | 211 | 0.48 |  |
|  |  | GN; NN | 317 | 0.52 |  |
|  |  | GG; GN | n/a |  | -- |
|  |  | NN; GN | 43 |  | 0.54 |
|  |  | GN; GN | 320 | 0.58* | 0.66**** |
| G | IL-N | GN; GG | 113 | 0.46 |  |
|  |  | GN; NN | 85 | 0.71** |  |
|  |  | GG; GN | n/a |  | -- |
|  |  | NN; GN | 250 |  | 0.53 |
|  |  | GN; GN | 104 | 0.37* | 0.64* |
| N | IL-N | GN; GG | 177 | 0.51 |  |
|  |  | GN; NN | 194 | 0.72**** |  |
|  |  | GG; GN | n/a |  | -- |
|  |  | NN; GN | 188 |  | 0.57 |
|  |  | GN; GN | 212 | 0.42 | 0.61* |

<sup>1</sup>Backcrosses using ILs (*M. guttatus* background = IL-G; *M. nasutus* background = IL-N) to the IM62 line of *M. guttatus* (G) and the SF5 line of *M. nasutus* (N). ♀ indicates the maternal parent and ♂ indicates the paternal parent.
<sup>2</sup>Two-locus genotype for *hms1* and *hms2*. GG = *M. guttatus* homozygote; GN = heterozygote; NN = *M. nasutus* homozygote.
<sup>3</sup>Number of progeny assessed. Two crosses were unsuccessful (labeled “no seeds”) because the IL-G with the genotype *hms1*<sub>GN</sub>; *hms2*<sub>NN</sub> was completely male sterile. The IL-N with the genotype *hms1*<sub>GG</sub>; *hms2*<sub>GN</sub> could not be generated (see text) and is labeled “n/a.”
<sup>4</sup>Percent *M. guttatus* (G) alleles at *hms1* transmitted to progeny from heterozygous IL parent.
<sup>5</sup>Percent *M. guttatus* (G) alleles at *hms2* transmitted to progeny from heterozygous IL parent.
\* $P < 0.05$ , \*\* $P < 0.01$ , \*\*\* $P < 0.005$ , \*\*\*\* $P < 0.0001$ based on $\chi^2$ tests of observed frequencies versus the Mendelian expectation.

The pattern of TRD at *hms2* follows what is expected if hybrid sterility acts through gametes. For example, if pollen grains are inviable when they carry *M. guttatus* alleles at *hms1* in combination with *M. nasutus* alleles at *hms2*, the effect of *hms1* paternal genotype on TRD at *hms2* should be additive. Indeed, progeny from males that carry one or two *M. guttatus* alleles at *hms1* show a 28% or 76% under-transmission of *M. nasutus* alleles at *hms2* relative to the Mendelian expectation (Figure S1). Consistent with the action of a gametic incompatibility, backcross progeny of doubly heterozygous IL parents (*i.e*., *hms1*_GN_; *hms2*_GN_) are much less likely to come from gametes with an *M. guttatus* allele at *hms1* in combination with an *M. nasutus* allele at *hms2* (Table 4). In these crosses, the *hms1*_G_; *hms2*_N_ gamete type is under-transmitted through both sexes, though the effect is stronger through males. Under-transmission is also more severe in crosses to IM62 (*M. guttatus*) and against the IL-N genetic background (Table S2).

**Table 4.**
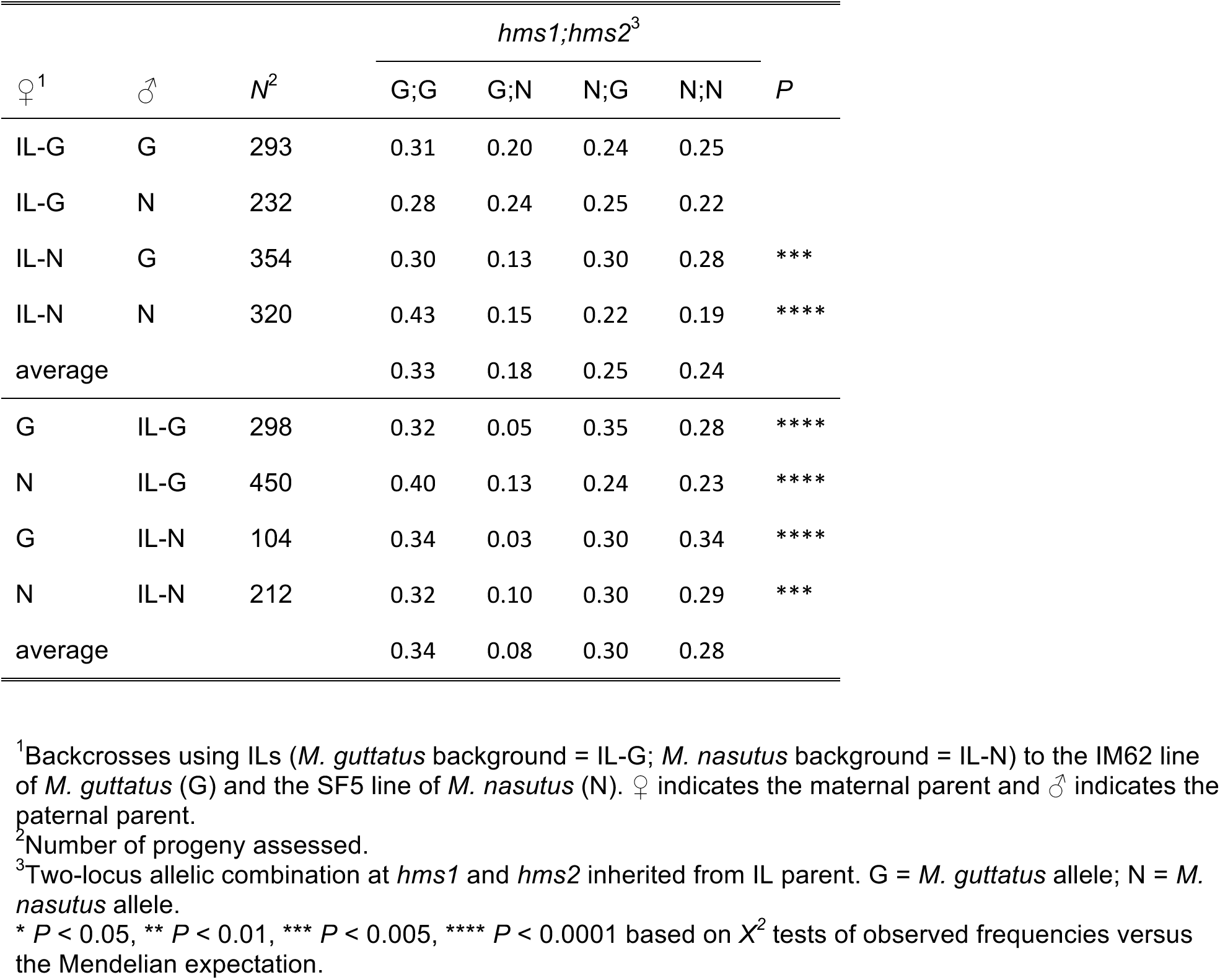
Two-locus transmission ratios at *hmsl* and *hms2* in backcross progeny of doubly heterozygous ILs.

If the *hms1-hms2* incompatibility acts through gametes, we might expect patterns of pollen viability to predict rates of transmission ratio distortion through males. To examine this possibility, we measured pollen viability in various two-locus genotypes of the IL-G and IL-Ns (Table 5). In general, patterns of male fertility and transmission ratio distortion are indeed related. For example, pollen viability is 64% in IL-Gs that are *hms1*_GG_; *hms2*_GN_. For this genotype, if we assume equal transmission of *M. guttatus* and *M. nasutus* alleles into pollen and attribute all sterility to *hms1*_G_; *hms2*_N_, then the *M. guttatus* allele at *hms2* should be present in 78% of progeny when this individual is used as the paternal parent in a cross (which is close to the observed frequency of 86%, Table 3). Similarly, for IL-Gs that are *hms1*_GN_; *hms2*_GN_, if we assume that all *hms1*_G_; *hms2*_N_ gametes are inviable (and divide the remaining 7% sterility equally among the other three two-locus genotypes), we expect *M. guttatus* allele frequencies of 33% and 66% at *hms1* and *hms2*, respectively. These values are very similar to what we observe when this IL-G genotype is backcrossed to *M. guttatus* (37% and 67%, Table 3).

**Table 5.** Pollen viability for various *hms1-2* IL genotypes.

| Genetic Background | *hms1; hms2* | *N*<sup>1</sup> | PV<sup>2</sup> |
| --- | --- | --- | --- |
| IL-G | GG; GN | 5 | 0.64 (0.04) |
|  | NN; GN | 16 | 0.79 (0.04) |
|  | GN; GN | 16 | 0.67 (0.06) |
|  | GN; GG | 12 | 0.71 (0.06) |
|  | GN; NN | 3 | 0.18 (0.17) |
| IL-N | NN; GN | 15 | 0.88 (0.02) |
|  | GN; GN | 14 | 0.81 (0.03) |
|  | GN; GG | 13 | 0.85 (0.02) |
|  | GN; NN | 18 | 0.09 (0.01) |

<sup>1</sup>Number of individuals scored.
<sup>2</sup>Pollen viability given as the proportion viable pollen grains per flower (for a haphazard sample of 100). PV is the average of two flowers and the number in parentheses is the standard error.

At *hms1*, TRD is more complex. On the one hand, *M. guttatus* alleles at *hms1* are under-transmitted due to the *hms1*_G_; *hms2*_N_ gametic sterility discussed above (Table S2). On the other hand, in many of the IL-backcrosses, *M. guttatus* alleles at *hms1* are overrepresented among the progeny (Tables 2 and 2.1). This effect is most pronounced when the IL parent is heterozygous at *hms1* and homozygous for *M. nasutus* alleles at *hms2* (Figure S1; note that this genotype is not completely sterile so crosses can still be performed). Remarkably, this direction of TRD is exactly the opposite of what is expected if *hms1* transmission is primarily influenced by the *hms1*_G_; *hms2*_N_ gametic incompatibility. Moreover, pollen viability in IL-G and IL-Ns with the genotype *hms1*_GN_; *hms2*_NN_ is much lower than the 50% expected for gametic expression of hybrid male sterility (Table 5), consistent with over-transmission of *M. guttatus hms1* alleles into pollen. Note that if these two TRD mechanisms – *hms1*_G_; *hms2*_N_ gamete sterility and over-transmission of *M. guttatus hms1* alleles – counteract each other in F_1_ hybrids and in doubly heterozygous ILs, it could explain why their progeny carry *hms1* alleles in roughly Mendelian proportions (Table 2, Figure S1). Consistent with this idea, backcross progeny of doubly heterozygous ILs are most often products of the *hms1*_G_; *hms2*_G_ gamete type (Table 4).

Additionally, a genetically distinct hybrid incompatibility appears to affect transmission of *hms1* against an *M. nasutus* genetic background. Self-fertilization of a doubly heterozygous IL-N individual produces no *M. guttatus* homozygotes at the *hms1* locus (Table 2), a genotype expected to appear in a quarter of the progeny (IL-N F_2_ *N* = 200, expected frequency = 50). When instead this same doubly heterozygous IL-N genotype is crossed to IM62 (in either direction), progeny homozygous for *M. guttatus* alleles at *hms1* are recovered (Table S3). Note that selfing the doubly heterozygous IL-N produces offspring with isogenic *M. nasutus* genetic backgrounds, whereas the backcross to IM62 results in progeny with genetic backgrounds that are F_1_-like. Taken together, these results suggest that the *hms1* region is involved in yet another hybrid incompatibility. This one causes lethality in hybrids that are homozygous for *M. guttatus* alleles at *hms1* (or linked loci) and homozygous for *M. nasutus* alleles at one or more unlinked loci.

By scoring genotype frequencies in the progeny of reciprocal backcrosses involving the doubly heterozygous ILs (*hms1*_GN_; *hms2*_GN_), it is possible to track which two-locus *hms1-2* meiotic products are transmitted through pollen and ovules. If we use these observed two-locus gametic allele frequencies (instead of assuming equal proportions of the four two-locus gamete types) to calculate expected genotype frequencies in the selfed progeny of doubly heterozygous ILs (i.e., IL-F_2_ populations), the resulting values do not significantly differ from observed proportions (Table 2, Table 4). To fully account for observed genotype frequencies in the IL-N F_2_, it is also necessary to assume complete lethality of *M. guttatus* homozygotes at *hms1* (Table 2; note that this hybrid lethality is not reflected in IL backcross allele frequencies because progeny do not carry the requisite *M. nasutus* genetic background for expression of the incompatibility).

In summary, we have identified at least three sources of *hms1-hms2* TRD in *M. nasutus-M. guttatus* F_2_ hybrids: 1) under-transmission of pollen and, to a lesser extent, ovules that carry an *M. guttatus* allele at *hms1* in combination with an *M. nasutus* allele at *hms2*, presumably due to gametic inviability, 2) over-transmission of *M. guttatus* alleles at *hms1*, an effect that occurs through males and females, and does not depend on genetic background, and 3) hybrid lethality in individuals homozygous for *M. guttatus* alleles at *hms1* (and linked genomic regions) in combination with *M. nasutus* homozygosity at one or more unlinked loci.

### Fine-mapping TRD

In previous (Sweigart and Flagel 2015) and ongoing efforts to fine-map *hms1* and *hms2*, we identified a small subset of SF5-IM62 F_2_ hybrids that were recombinant for one or both sets of *hms* flanking markers. With the goal of genetically mapping TRD in both regions, we self-fertilized these recombinants to generate F_3_ progeny and examined genotype frequencies at both sets of flanking markers (Figures 3 and 4). We reasoned that TRD in the F_3_ progeny should only be observable if the causal locus is heterozygous in the F_2_ parent. If, instead, the TRD-causing locus is homozygous (for either *M. guttatus* or M. *nasutus* alleles), loci in the adjacent heterozygous region should segregate in a Mendelian fashion.

As in the IL crosses, patterns of *hms2*-linked TRD were consistent with the action of *hms1*_G_; *hms2*_N_ gametic sterility. In this genomic region, the most extreme TRD occurred in the two F_3_ families that descended from F_2_ hybrids with the *hms1*_GG_; *hms2*_GN_ genotype (Figure 2). Despite this general support for *hms1-hms2* gametic sterility, *hms2*-linked TRD could not be unambiguously mapped to a particular genomic region (no interval in Figure 2 is perfectly associated with presence/absence of TRD). Presumably, genetic background in these F_2_ hybrids can mask TRD associated with *hms1*_G_; *hms2*_N_ gametic sterility (*e.g*., 28_22) or mimic it (*e.g*., 02_66).

**Figure 2.**
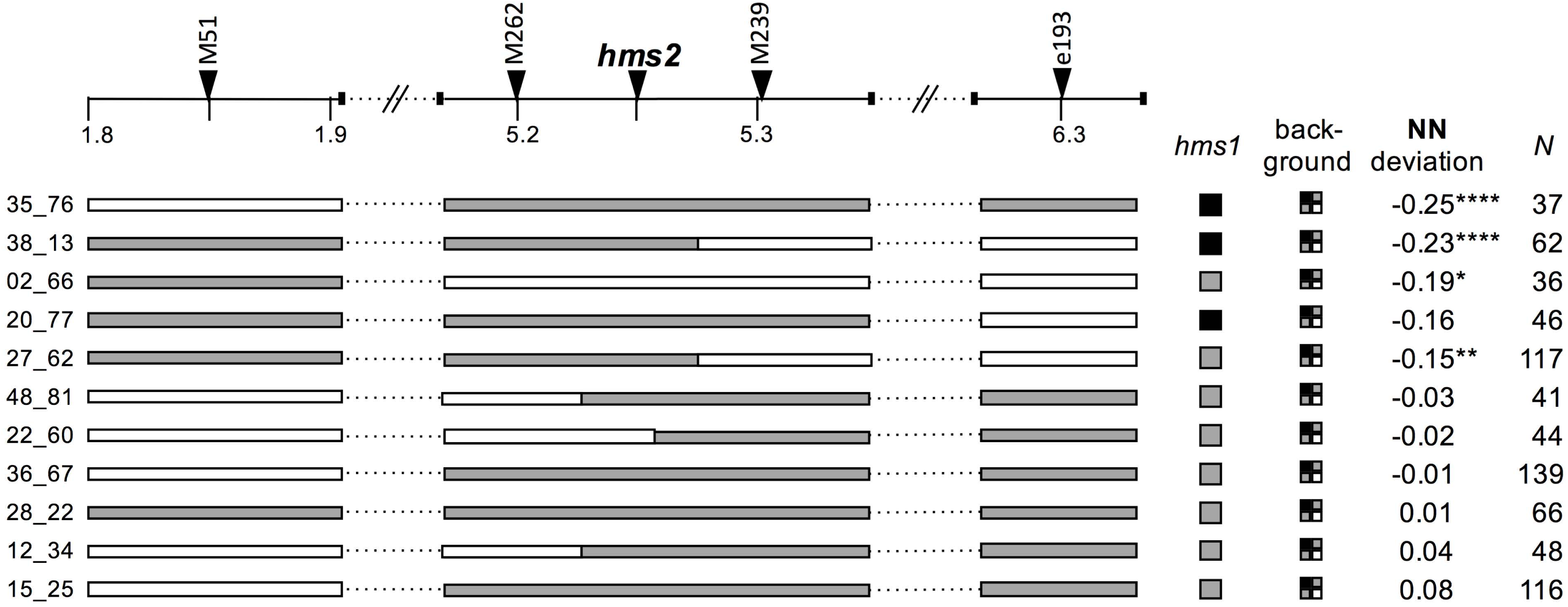
Genetic dissection of /ww2-linked TRD in *Mimulus.* A physical map of ~4.5 Mb the *hms2* region is shown, including the positions of genetic markers (indicated with triangles along the top). F_2_ recombinants are shown with horizontal bars representing genotypes in the genomic region linked to *hms2* and squares indicating genotypes at *hmsl* and across the genetic background (white = *M. nasutus* homozygote, grey = heterozygote, black = *M. guttatus* homozygote). Deviation from the Mendelian expectation (0.25) ofM *nasutus* homozygotes (NN)in the F_3_ progeny is given. *N*indicates the number of F_3_ progeny scored from each individual. * *P <* 0.05, ** *P <* 0.01, *** *P <* 0.005, **** *p <* 0.0001 based onX^2^ tests of observed frequencies versus the Mendelian expectation.

At *hms1*, the two contributors to TRD were decoupled in F_2_ recombinants with *M. guttatus* homozygotes overrepresented in some F_3_ families and underrepresented in others (Figure 3). As with the IL experiments, the most significant over-transmission of *M. guttatus* alleles at *hms1* appears in the progeny of F_2_ hybrids that are homozygous for *M. nasutus* alleles at *hms2* (Figure 3, first two F_2_s). This TRD phenotype maps to an 800-kb region that includes *hms1*, but we have too few recombinants to determine if the hybrid TRD phenotype is genetically separable from hybrid sterility. For a distinct set of *hms1* F_2_ recombinants, we observed a severe deficit of *M. guttatus* homozygotes among their F_3_ progeny (Figure 3, last six F_2_ individuals), consistent with the expression of hybrid lethality as seen in the IL experiments. This TRD phenotype maps to at least two independent loci in the *hms1* region and is not affected by *hms2* genotype, suggesting a distinct genetic basis for this hybrid incompatibility.

**Figure 3.**
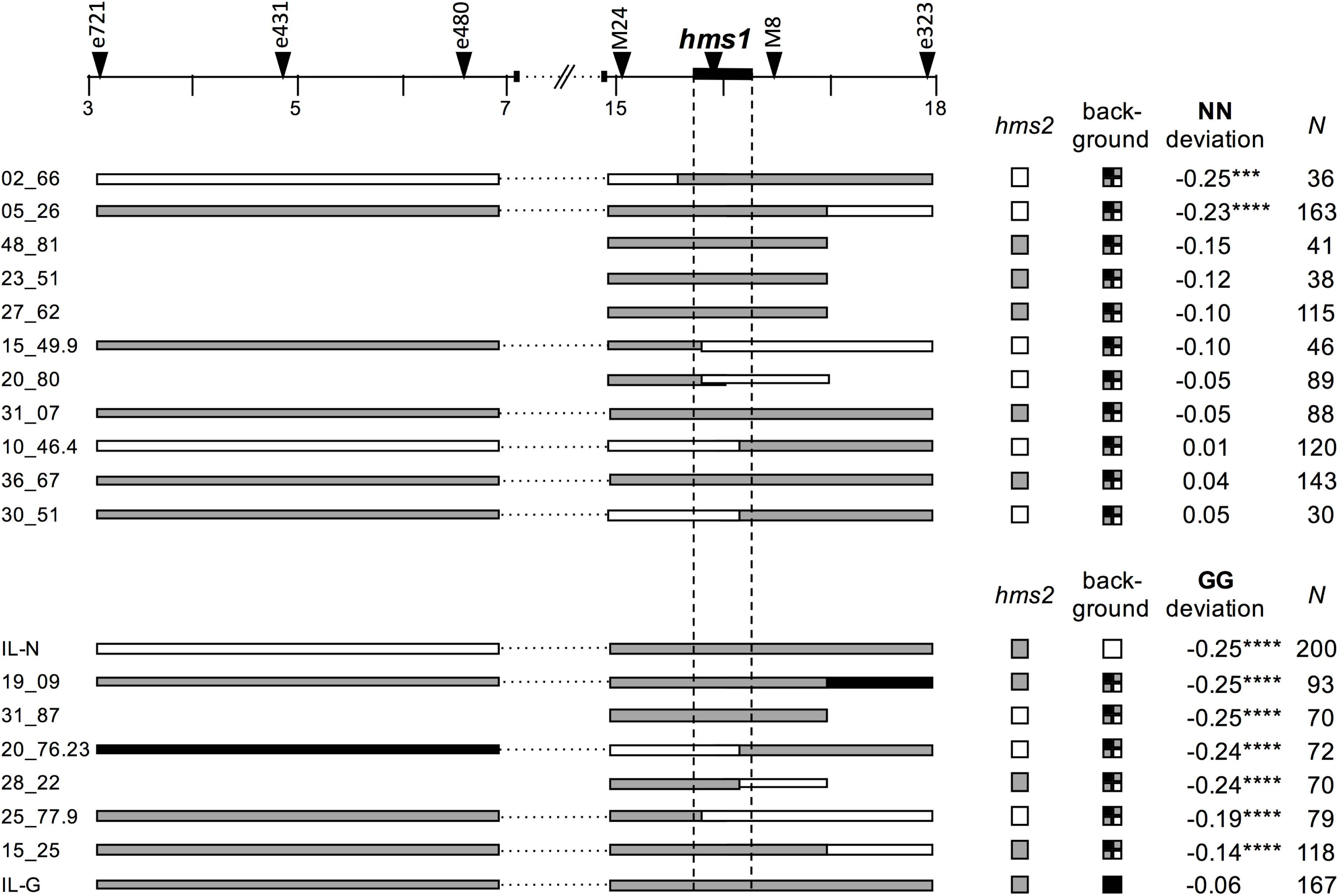
Genetic dissection of *hmsl*-linked TRD in *Mimulus.* A physical map of 15 Mb the *hmsl* region is shown, including the positions of genetic markers (indicated with triangles along the top) and the 320-kb *hmsl* haplotype (shown as a solid black bar with dotted lines extending downwards). F_2_ recombinants are shown with horizontal bars representing genotypes in the genomic region linked to *hmsl* and squares indicating genotypes at *hms2* and across the genetic background (white = *M. nasutus* homozygote, grey = heterozygote, black = *M. guttatus* homozygote). Deviation from the Mendelian expectation (0.25) of *M. nasutus* homozygotes (NN) in the F_3_ progeny is given for the top group of 11 F_2_ recombinants. Deviation from the Mendelian expectation (0.25) of *M. guttatus* homozygotes (GG) in the F_3_ progeny is given for the bottom group of six F_2_ recombinants and the doubly heterozygous ILs. *N* indicates the number of F_3_ progeny scored from each individual. *** *P* < 0.005, **** *P* < 0.0001 based on *X^2^* tests of observed frequencies varsus the Mendelian expectation.

### TRD at *hms1* within *M. guttatus*

To investigate whether *hms1*-linked TRD is a strictly hybrid phenomenon or also occurs within *M. guttatus*, we generated reciprocal F_2_ progeny between IM62 and IM767. These two inbred lines carry distinct alleles at *hms1* and show very different patterns of variation in the surrounding genomic region. The IM62 line carries an incompatible, hybrid sterility-causing *hms1* allele embedded within a distinctive, 320-kb haplotype, whereas IM767 carries a compatible (*i.e*., non-sterility causing) allele at *hms1* and typical levels of nucleotide variation in the region (Sweigart and Flagel 2015). Because genotype frequencies at *hms1* did not differ significantly between reciprocal F_2_ populations (data not shown), we pooled data from both directions of the cross. We observed modest, but significant TRD at *hms1* with an excess of IM62 homozygotes (frequency of IM62 homozygotes to heterozygotes to IM767 homozygotes: expected 0.25:0.5:0.25, observed 00.27:0.54:0.19, *X^2^* = 6.479, d.f. = 2, *P* = 0.0027, *N* = 582). However, the bias in allelic transmission toward IM62 was not significant (frequency of IM62:IM767 alleles: expected 0.5:0.5, observed 0.54:0.46, *X^2^* = 0.151, d.f. = 1, *P* < 0.151, *N* = 582) and genotype frequencies did not significantly differ from the expectation given the allele frequencies (*X^2^* = 2.025, d.f. = 2, *P* = 2.025, *N* = 582). To further investigate the mechanism of *hms1*-linked TRD, we performed reciprocal backcrosses using IM62 and IM767. However, unlike in the IM62-IM767 F_2_ hybrids, all four backcross populations exhibited nearly perfect Mendelian ratios (expected 0.50:0.50; F_1_ x IM62 = 0.50:0.50, *N* = 279; F_1_ x IM767 = 0.50:0.50, *N* = 281; IM62 x F_1_ = 0.51:0.49, *N* = 189; IM767 x F_1_ = 0.49:0.51, *N* = 188). These results suggest there is little to no transmission bias favoring the *hms1* incompatibility allele or the associated 320-kb haplotype within the Iron Mountain population.

Finally, we wanted to investigate if the presence of *M. nasutus* alleles at *hms2* increases the transmission bias of IM62 at *hms1* – even in an otherwise *M. guttatus* genetic background. To address this question, we examined genotype frequencies in the reciprocal backcross progeny of individuals that were heterozygous IM62/IM767 at *hms1* and segregating for an *M. nasutus* introgression at *hms2* (against an otherwise IM62-IM767 F_2_ genetic background; Figure S2). Indeed, extreme TRD at *hms1* (*i.e*., bias toward the IM62 allele > 70%) was only observed in the backcross progeny of one individual (08_60) that was also homozygous for *M. nasutus* alleles at *hms2* (Table 6). These results suggest that over-transmission of the IM62 allele at *hms1*, which appears to require *M. nasutus* alleles at *hms2*, may occur exclusively in hybrids.

**Table 6.** Transmission of IM62 vs. IM767 at *hms1* varies depending on *hms2* genotype.

| *hms2*<br>genotype | F<sub>2</sub> ID<sup>1</sup> | %IM62<sup>2</sup> |  |
| --- | --- | --- | --- |
|  |  | F<sub>2</sub> male | F<sub>2</sub> female |
| IM62 | 02_02 | 0.58 (74) | 0.55 (64) |
|  | 02_46 | 0.48 (121) | 0.43 (28) |
|  | 06_31 | 0.55 (179) | 0.29 (41) |
|  | 06_70 | 0.55 (123) | 0.50 (116) |
|  | 06_96 | 0.41 (46) | -- |
|  | combined | 0.53 (543) | 0.47 (249) |
| IM767 | 02_17 | 0.45 (53) | 0.56 (122) |
|  | 02_48 | 0.54 (79) | 0.49 (84) |
|  | 02_68 | 0.56 (39) | 0.49 (141) |
|  | 06_39 | 0.55 (107) | -- |
|  | combined | 0.53 (278) | 0.51 (347) |
| SF | 08_60 | 0.77 (104)**** | 0.73 (75)*** |
|  | 12_09 | 0.50 (111) | 0.54 (41) |
|  | combined | 0.62 (215)** | 0.66 (116)* |

<sup>1</sup>Individual IDs for F<sub>2</sub> progeny from BG<sub>4</sub>275-IM767 crosses. At *hms1*, all F<sub>2</sub> individuals used were heterozygous for IM62 and IM767 alleles; at *hms2*, individuals used were homozygous for IM62, IM767, or SF alleles (see text for details).
<sup>2</sup>Percent IM62 alleles at *hms1* transmitted to progeny from IM62-IM767 heterozygous parent. Value given in parentheses is the number of progeny assessed.
\* $P < 0.05$ , \*\* $P < 0.01$ , \*\*\* $P < 0.005$ , \*\*\*\* $P < 0.0001$ based on $\chi^2$ tests of observed genotype frequencies versus the Mendelian expectation.

## DISCUSSION

Transmission ratio distortion is commonly observed among hybrid offspring of recently diverged species, but the evolutionary significance is not always clear. In this study, we identified multiple contributors to hybrid TRD in genomic regions linked to two *Mimulus* hybrid sterility loci *hms1* and *hms2*, revealing a fine-scale complexity reminiscent of several previously characterized hybrid incompatibilities (Davis and Wu 1996; Long et al. 2008; Yang et al. 2012; Kubo et al. 2016b). We have discovered that hybrid transmission bias is caused, in part, by gametic action of the *hms1-hms2* incompatibility itself. However, the effects of the gametic hybrid sterility are partially obscured by an opposing (and currently unknown) mechanism that results in over-transmission of the *M. guttatus hms1* incompatibility allele in certain hybrid genetic backgrounds. In addition, our genetic analyses uncovered an independent hybrid lethality system with at least two incompatibility loci tightly linked to *hms1*. Strikingly, we found no evidence of biased transmission of the *hms1* incompatibility allele within *M. guttatus*, providing little support for selfish evolution as the cause of a recent, partial sweep at *hms1* (Sweigart and Flagel 2015). Instead, it appears that TRD at *hms1* and *hms2* might occur exclusively in hybrids.

### Gametic action of *hms1-hms2* hybrid incompatibility

Our finding that the *hms1*_G_; *hms2*_N_ gamete type is severely under-transmitted in six of the eight backcrosses involving doubly heterozygous ILs (*hms1*_GN_; *hms2*_GN_) is strong evidence of gametic action of the incompatibility. This result runs counter to our previous interpretation of the finding that pollen viability is reduced from the F_1_ to F_2_ generation, which seemed to suggest a diploid (sporophytic) genetic basis for the *hms1-hms2* incompatibility (Sweigart et al. 2006). In general, for a hybrid incompatibility that affects the gametophyte, sterility is expected to be less severe in the F_2_ generation due to the inviability of recombinant F_1_ gametes and regeneration of parental combinations. However, in this case, it appears that removal of *hms1*_G_; *hms2*_N_ F_1_ gametes is somewhat balanced by over-transmission of *M. guttatus* alleles at *hms1*. Moreover, incomplete penetrance of F_1_ hybrid gametic sterility (*i.e*., some *hms1*_G_; *hms2*_N_ gametes do contribute to the F_2_ generation, see Table 4) produces a small fraction of F_2_ hybrids that are completely sterile because they are homozygous for incompatible alleles (*i.e*., *hms1*_GG_; *hms2*_NN_).

As an independent line of evidence for gametic expression of the *hms1-hms2* incompatibility, it is apparently difficult to introgress *M. nasutus hms2* alleles into an *M. guttatus* genetic background. In the BG_4_-NIL population (generated by four rounds of backcrossing using IM62 as the pollen donor; see Methods from this study and Fishman and Willis 2005), only 2.8% of individuals (5/175) are heterozygous at MgSTS45, a marker ~2 cM from *hms2* (L. Fishman, unpublished results). This level of distortion is notable: of the 194 markers genotyped in this BG_4_ population, only four of them show lower heterozygosity and three of those map near a meiotic drive locus that strongly favors the *M. guttatus* allele (Fishman and Saunders 2008). In the BN_4_ population (generated by four rounds of backcrossing with SF5 as the pollen donor), heterozygous introgressions at MgSTS45 are much more common, occurring in 10% of individuals (18 of 181). This result is not unexpected given that *M. guttatus* alleles at *hms2* are perfectly compatible with *M. nasutus* alleles at *hms1*.

Unlike in animals, hybrid incompatibilities in plants are often gametic (Morishima et al. 1991; Koide et al. 2008b; Leppala et al. 2013). Based on his studies of hybrid sterility between the *indica* and *japonica* varieties of *Oryza sativa*, Oka (1974) first suggested that defects in pollen development might be caused by loss-of-function alleles at duplicate genes (Oka 1974). Indeed, two cases of this duplicate gametic lethal model have now been demonstrated at the molecular level (Mizuta et al. 2010; Yamagata et al. 2010). For *Mimulus hms1* and *hms2*, there’s no evidence that gene duplicates are involved (Sweigart and Flagel 2015), but a similar pattern of hybrid sterility is expected to result from a two-locus hybrid incompatibility between any genes expressed in the gametophyte. Additionally, the fact that the *hms1-hms2* incompatibility seems to affect both the male and female gametophyte (the *hms1*_G_; *hms2*_N_ gamete type is under-transmitted through both sexes) is consistent with our finding that these loci contribute to both hybrid male sterility and hybrid female sterility (Sweigart et al. 2006). Gametic hybrid incompatibilities that affect the fertility of both sexes have also been discovered in tomato, rice, and *Arabidopsis* (Rick 1966; Koide et al. 2008a; Leppala et al. 2013), though they are apparently less common than those that act in only one sex (Morishima et al. 1991; Koide et al. 2008b)

### Additional sources of transmission ratio distortion

Our fine-scale dissection of TRD at *hms1* and *hms2* provides insight into genomic differentiation between closely related *Mimulus* species and reveals a potentially complex genetic basis for hybrid dysfunction. In other systems, fine-mapping has often revealed multiple, tightly linked hybrid incompatibility loci that show independent effects (Wu and Davis 1993; Kubo et al. 2016a; Simon et al. 2016) or epistasis (Long et al. 2008; Yang et al. 2012; Kubo et al. 2016b). In one particularly complex example from *indica* and *japonica*, fine-mapping revealed two tightly linked genes involved in independent two-locus pollen killer systems (Kubo et al. 2016b). Because of this tight linkage, pollen killing had initially appeared to be caused by a single, three-locus interaction (Kubo *et al*. 2008). Remarkably, both of these pollen killer systems involve interactions between sporophytic and gametophytic genes, as well as additional modifier loci (Kubo et al. 2016b). The picture emerging from such studies is one of hybrid sterility regulated by multiple, interconnected molecular networks, potentially involving many genes.

A key question for *hms1* and *hms2* is whether the same genes cause the gametic incompatibility and transmission bias of *M. guttatus* at *hms1*. The latter is particularly strong when *hms2* is homozygous for *M. nasutus* alleles (Table 3, Figure S1), suggesting it might be caused by an interaction between the two loci. Additionally, the presence of *hms2*_NN_ also appeared to unleash severe *hms1* TRD in one of the two IM62-IM767 F_2_ populations in which it was present (Table 6), suggesting *hms2* might be necessary but not sufficient for *hms1* TRD. On the other hand, over-transmission of *hms1*_G_ does not seem to absolutely require *hms2*_NN_ (*e.g*., we observed 62% transmission of *hms1*_G_ in *M. nasutus* x IL-G_GN;GG_, Table 3), which might argue against its direct involvement. Indeed, for the IL-Gs, there is a bias toward *hms1*_G_ in all backcross populations except those involving doubly heterozygous IL parents (*i.e*., *hms1*_GN_; *hms2*_GN_), which, because they express the *hms1*_G_; *hms2*_N_ gametic inviability, might obscure additional sources of *hms1* TRD. Going forward, additional rounds of high-resolution fine-mapping will be needed to pinpoint the causal genes and determine if *Mimulus* hybrid sterility and TRD are genetically separable. Such efforts in rice have been successful in disentangling the complex phenotypic effects of linked hybrid sterility loci (*e.g*., (Kubo et al. 2016a).

Identifying the molecular genetic basis of *hms1* TRD might also provide insight into its mechanisms. Because the bias toward *M. guttatus* alleles at *hms1* occurs through both males and females, the simplest single explanation is a gamete killing system that affects pollen and seeds. Alternatively, it is possible that independent mechanisms (and genetic loci) cause sex-specific TRD, such as pollen competition in males (*e.g*., (Fishman et al. 2008)) and meiotic drive in females (*e.g*., (Fishman and Saunders 2008). Whatever the cause, over-transmission of *hms1*_G_ is apparently exacerbated by *M. nasutus* alleles at *hms2* to the point of overwhelming the effects of the *hms1*_G_; *hms2*_N_ gametic incompatibility. Indeed, the direction of TRD in the backcross progeny of *hms1*_GN_; *hms2*_NN_ ILs is counterintuitive: because of the *hms1*_G_; *hms2*_N_ gametic incompatibility, one expects transmission bias to be toward *M. nasutus* alleles. Instead, we observed exactly the opposite, namely, strong transmission bias toward *M. guttatus* at *hms1*. This finding might help explain < 50% pollen inviability in ILs with the genotype *hms1*_GN_; *hms2*_NN_. If *hms1*_G_ alleles are highly overrepresented in pollen of such individuals due to gamete killing or some other mechanism, the gametic incompatibility will be expressed more often than expected under Mendelian inheritance. However, to explain the bias toward *M. guttatus* alleles in the backcross progeny, the gamete killing phenotype has to be stronger than the gametic incompatibility. In other words, some fraction of *hms1*_G_; *hms2*_N_ gametes must survive – and in greater numbers than *hms1*_N_; *hms2*_N_ gametes – to form zygotes. Clarifying the role of *hms2* in *hms1* TRD, and whether it acts through the diploid sporophyte or haploid gametophyte, will be an important step toward understanding the mechanistic basis of hybrid distortion.

Surprisingly, our crossing experiments revealed at least two additional hybrid incompatibility loci linked to *hms1*. These loci, which contribute to TRD in the IL-Ns, appear to cause hybrid inviability and involve recessive alleles from both *Mimulus* species: against an *M. nasutus* genetic background, the *hms1* region cannot be made homozygous for *M. guttatus* alleles. The precise locations of these hybrid lethality loci are not yet known (Figure 3), but both potentially overlap with the 320-kb haplotype associated with the *hms1* incompatibility allele (Sweigart and Flagel 2015). This nearly invariant haplotype, which includes 30 genes, has recently risen to intermediate frequency in the Iron Mountain population of *M. guttatus*. The fact that multiple hybrid incompatibility loci are associated with this sweeping haplotype suggest that natural selection within a single population might have profound consequences for reproductive isolation between *Mimulus* species.

### Implications for the evolution of hybrid sterility in *Mimulus*

An emerging theme in speciation genetics is that selfish evolution within species might be a major driver of hybrid incompatibilities. Decades of genetic analysis have provided a detailed mechanistic understanding of classic segregation distorters within *Drosophila* and mouse species (see (Presgraves 2008), and more recent studies have shown that hybrid sterility and hybrid TRD can be caused by the same genes (Phadnis and Orr 2009a; Zhang et al. 2015). However, very few studies have directly linked these two ends of the spectrum, testing whether incompatibility alleles act as selfish genetic elements within species. In one recent exception, Case *et al*. (2016) showed population genomic evidence for coevolution between a selfish cytoplasmic male sterility (CMS) gene and a nuclear restorer of fertility (*Rf* locus) within the Iron Mountain population of *M. guttatus* (Case et al. 2016). These same two loci also cause hybrid male sterility between *M. guttatus* and *M. nasutus*, suggesting that intragenomic conflict within Iron Mountain contributes to interspecific reproductive barriers.

Direct evidence for selfish evolution is missing from any of the hybrid gamete eliminators that have been cloned in rice (Long et al. 2008; Kubo et al. 2011; Yang et al. 2012; Kubo et al. 2016a; Kubo et al. 2016b; Yu et al. 2016). In most of these hybrid sterility systems, patterns of molecular variation at the causal genes in *japonica*, *indica*, and their wild ancestor *Oryza rufipogon* suggest that hybrid incompatibility alleles may never have expressed their killing phenotypes within species (*e.g*., (Long et al. 2008; Yang et al. 2012),; also see (Sweigart and Willis 2012). In plants, it is also important to consider that even if gamete eliminators do arise within species and evolve selfishly to bias their own transmission, they might do so without any cost to individual fitness (Rick 1966). Especially for pollen killers, a sufficient number of viable pollen grains might still remain to fertilize all available ovules. Under a scenario of selfish evolution with no fitness costs, there is no conflict and, thus, no mechanism for generating hybrid incompatibilities.

Despite evidence for a recent selective sweep of the *hms1*-associated haplotype in the Iron Mountain population (Sweigart and Flagel 2015), our crossing experiments suggest there is no transmission bias favoring the IM62 *hms1* incompatibility allele. One caveat to this finding is that TRD at *hms1* might vary in different genetic backgrounds; even if there is no transmission bias between the IM62 and IM767 *hms1* alleles, TRD might occur in other heterozygous combinations. Alternatively, Iron Mountain individuals, including IM62 and IM762, might carry suppressors at *hms2*. However, given the recentness of the *hms1*-associated sweep (i.e. ~63 generations old; Sweigart and Flagel 2015), it seems unlikely that there has been sufficient time for a suppressor to evolve. Instead, *M. guttatus* from Iron Mountain and elsewhere may carry a “permissive” allele at *hms2* that allowed the evolution of the IM62 *hms1* variant without it expressing any transmission bias or sterility. Consistent with this idea, the incompatibility allele at *hms2* seems to be specific to *M. nasutus* (Sweigart et al. 2007), indicating this species likely carries the derived allele. Thus, instead of being driven by selfish evolution within *M. guttatus*, it appears that TRD at *hms1* is limited only to hybrids. These findings leave open the possibility that *hms1* evolution within Iron Mountain may have been driven by ecological adaptation. Further molecular characterization of these hybrid incompatibility loci and direct investigations of the fitness effects of alternative alleles at *hms1* will be important steps toward identifying the evolutionary causes of this reproductive barrier.

## ACKNOWLEDGEMENTS

We thank Lila Fishman for sharing her BG_4_ and BN_4_ NILs and for valuable discussions. We are also grateful to Matt Zuellig who made thoughtful comments on an earlier draft, which improved the manuscript. We are especially indebted to Taylor Harrell and Rachel Hughes for expert greenhouse care and genotyping assistance. This work was supported by a National Science Foundation grant DEB-1350935 and funds from the University of Georgia Research Foundation to ALS.

---

**Figure S1.**
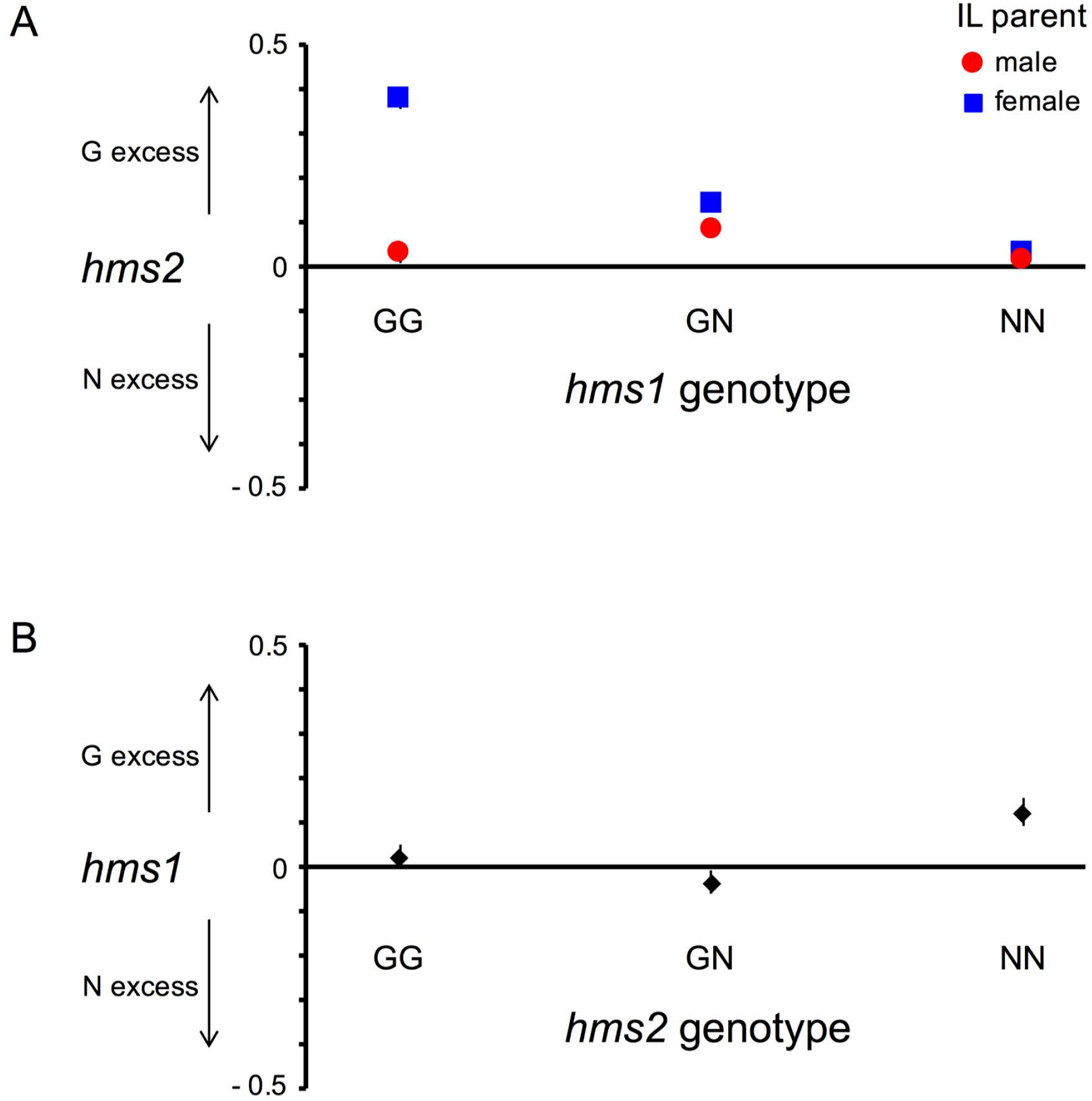
Transmission ratio distortion at *hmsl* and *hms2* in IL-backcross progeny. The vertical position of each symbol shows the magnitude and direction of the deviation of allelic transmission from the Mendelian expectation (0.5). *M. guttatus* deviations are graphed directly [deviation = *f*(N − 0.5)], and the *M. nasutus* deviations are graphed as negative [deviation = - (*f*(N-0.25)]. Thus, values above zero indicate excess ofM *guttatus* alleles and values below zero indicate excess of *M. nasutus* alleles. G = *M. guttatus* allele, N =*M. nasutus* allele. A) Allelic transmission of *hms2m* the progeny of IL-backcross es is significantly affected by IL parental genotype at *hmsl* (*F*= 37.6919, *P <* 0.0001), cross direction (*F*= 72.3339, *P<* 0.0001), and their interaction (*F* = 31.8353, *P <* 0.0001). B) Allelic transmission of *hmsl* in the progeny of IL-backcrosses is significantly affected by IL parental genotype at *hmsl* (*F* = 7.7977, *P* = 0.0043).

**Figure S2.**
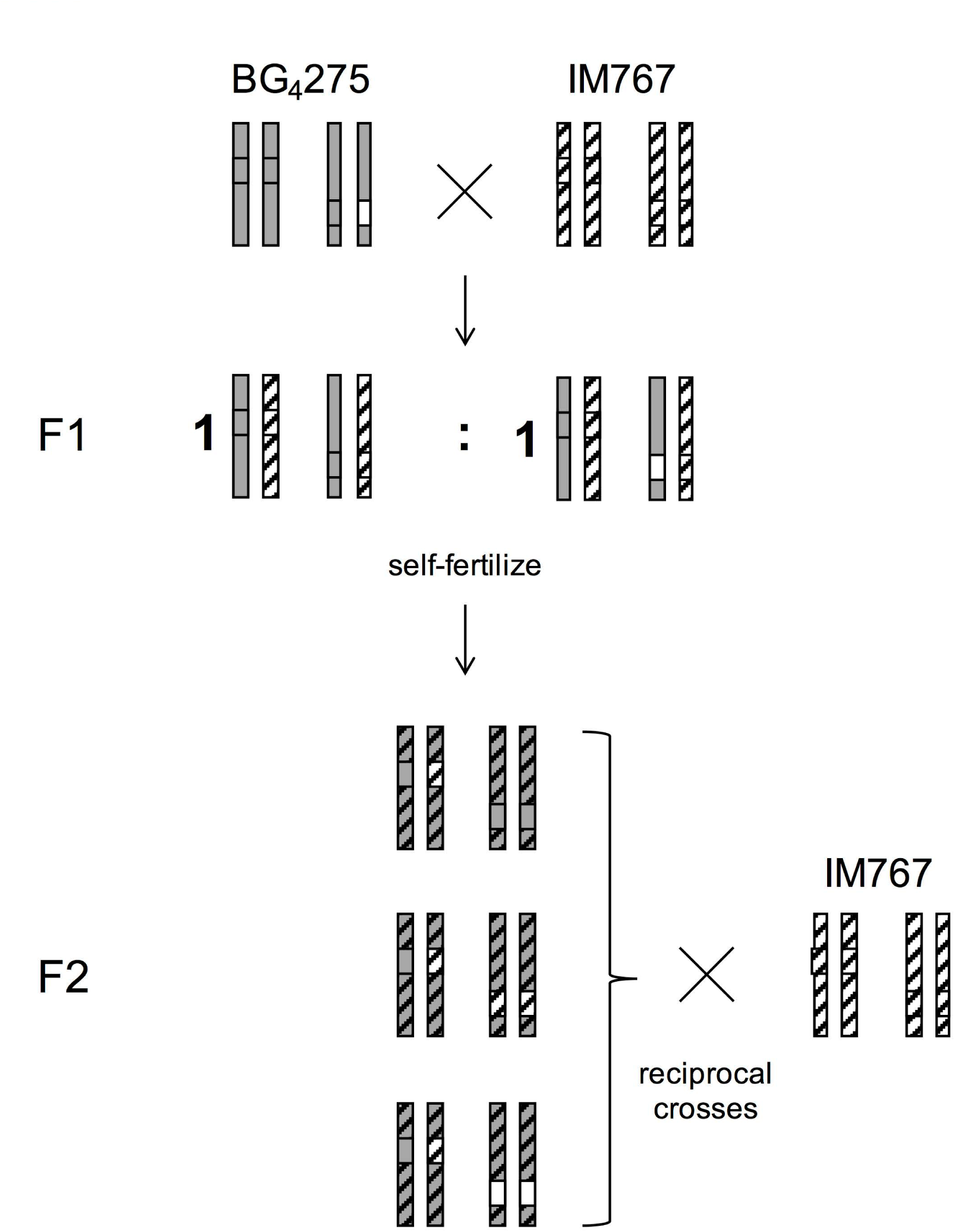
Crossing scheme for investigating the effect of *M*. *nasutus* (SF5) homozygosity at *hms2* on within-M *guttatus* TRD at *hmsl.* For each genotype, two chromosome pairs are shown (one with *hmsl* and one with *hms2*). We intercrossed IM767 (stripes) and BG_4_.275, which carries a heterozygous SF5 introgression (white) at *hms2* against an IM62 (grey shading) genetic background. The resulting progeny segregate 1:1 for two different genotypes at *hms2*; IM62-IM767 heterozygotes and SF5-IM767 heterozygotes (the remaining genetic background is heterozygous for IM62 and IM767 alleles). We genotyped F_2_ progeny with *hms*-linked markers to identify IM62-IM767 *hmsl* heterozygotes in combination with three different *hms2* genotypes: IM62 homozygotes, IM767 homozygotes, and SF5 homozygotes. Individuals with each of these three two-locus genotypes were then reciprocally backcrossed to IM767 to assess TRD at *hmsl.* Note that the genetic background of the F_2_ progeny are expected to segregate 1:2:1 for IM62 homozygotes, heterozygotes, and IM767 homozygotes (grey shading with stripes).

**Table S1.**
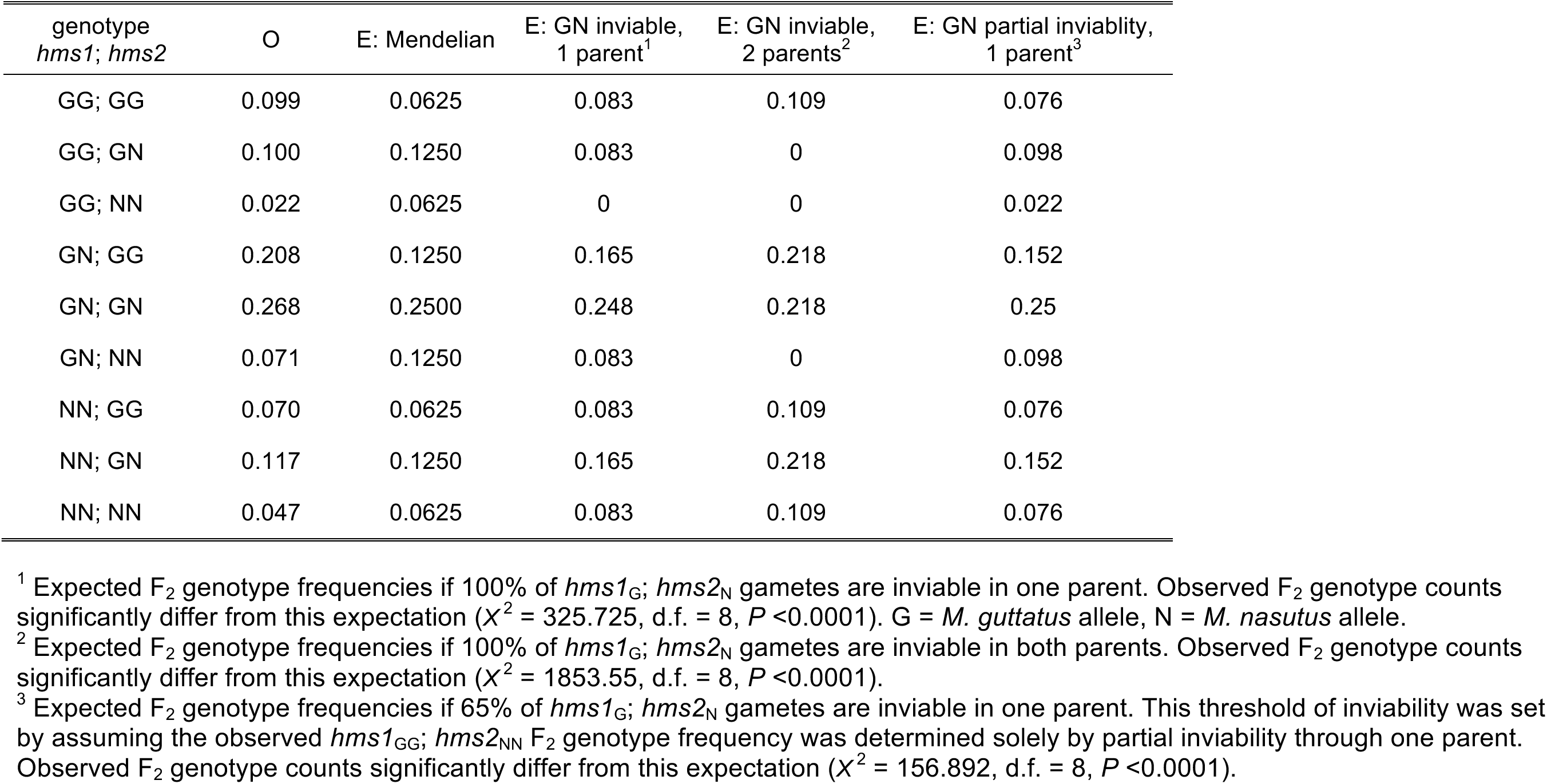
Observed and expected genotype frequencies at *hms1* and *hms2* in F_2_ hybrids *(N* = 5487).

| genotype<br>*hms1*; *hms2* | O | E: Mendelian | E: GN inviable,<br>1 parent<sup>1</sup> | E: GN inviable,<br>2 parents<sup>2</sup> | E: GN partial inviability,<br>1 parent<sup>3</sup> |
| --- | --- | --- | --- | --- | --- |
| GG; GG | 0.099 | 0.0625 | 0.083 | 0.109 | 0.076 |
| GG; GN | 0.100 | 0.1250 | 0.083 | 0 | 0.098 |
| GG; NN | 0.022 | 0.0625 | 0 | 0 | 0.022 |
| GN; GG | 0.208 | 0.1250 | 0.165 | 0.218 | 0.152 |
| GN; GN | 0.268 | 0.2500 | 0.248 | 0.218 | 0.25 |
| GN; NN | 0.071 | 0.1250 | 0.083 | 0 | 0.098 |
| NN; GG | 0.070 | 0.0625 | 0.083 | 0.109 | 0.076 |
| NN; GN | 0.117 | 0.1250 | 0.165 | 0.218 | 0.152 |
| NN; NN | 0.047 | 0.0625 | 0.083 | 0.109 | 0.076 |

<sup>1</sup> Expected F<sub>2</sub> genotype frequencies if 100% of *hms1*<sub>G</sub>; *hms2*<sub>N</sub> gametes are inviable in one parent. Observed F<sub>2</sub> genotype counts significantly differ from this expectation ( $\chi^2 = 325.725$ , d.f. = 8, $P < 0.0001$ ). G = *M. guttatus* allele, N = *M. nasutus* allele.
<sup>2</sup> Expected F<sub>2</sub> genotype frequencies if 100% of *hms1*<sub>G</sub>; *hms2*<sub>N</sub> gametes are inviable in both parents. Observed F<sub>2</sub> genotype counts significantly differ from this expectation ( $\chi^2 = 1853.55$ , d.f. = 8, $P < 0.0001$ ).
<sup>3</sup> Expected F<sub>2</sub> genotype frequencies if 65% of *hms1*<sub>G</sub>; *hms2*<sub>N</sub> gametes are inviable in one parent. This threshold of inviability was set by assuming the observed *hms1*<sub>GG</sub>; *hms2*<sub>NN</sub> F<sub>2</sub> genotype frequency was determined solely by partial inviability through one parent. Observed F<sub>2</sub> genotype counts significantly differ from this expectation ( $\chi^2 = 156.892$ , d.f. = 8, $P < 0.0001$ ).

**Table S2.** The severity of under-transmission of *hms1*_G_; *hms2*_N_ gametes (measured as the deviation from the Mendelian expectation of 0.25) in IL-backcrosses is affected by genetic background, cross direction, and identity of the recurrent parent.

| Effect<sup>1</sup> | df | *F* | *P* | LSM |  |
| --- | --- | --- | --- | --- | --- |
| Background<sup>2</sup> | 1 | 8.259 | 0.045 | G: -0.095 | N: -0.149 |
| Cross direction<sup>3</sup> | 1 | 30.910 | 0.005 | ♂: -0.174 | ♀: -0.070 |
| Recurrent parent<sup>4</sup> | 1 | 7.359 | 0.053 | G: -0.147 | N: -0.097 |

<sup>1</sup>Effects assessed by ANOVA with degrees of freedom (df), F-ratio (*F*), p-values (*P*), and least squares means (LSM) indicated.
<sup>2</sup>Crosses performed using fourth-generation NILs. *M. nasutus* background = BN<sub>4</sub>; *M. guttatus* background = BG<sub>4</sub>.
<sup>3</sup>Cross direction indicates whether the NIL was used as the paternal (♂) or maternal (♀) parent.
<sup>4</sup>Testcrosses were to the IM62 line of *M. guttatus* (G) or the SF line of *M. nasutus* (N).

**Table S3.** Genotype counts for progeny from reciprocal backcrosses between IM62 and the doubly heterozygous IL-N (*hms1*_GN_; *hms2*_GN_).

| Maternal parent | *hms1*; *hms2* genotype |  |  |  |
| --- | --- | --- | --- | --- |
|  | GG; GG | GG; GN | GN; GG | GN; GN |
| IL-N | 105 | 45 | 105 | 99 |
| IM62 | 35 | 3 | 31 | 35 |

